# Neophobia and innovation in critically endangered Bali myna, *Leucopsar rothschildi*

**DOI:** 10.1101/2021.11.12.468403

**Authors:** Rachael Miller, Elias Garcia-Pelegrin, Emily Danby

## Abstract

Cognition underlies animal behaviour, which is key to successful conservation strategies, yet largely under-utilised in conservation, though there are recent calls for closer integration. Conservation-relevant cognitive abilities can impact on adaptability and survival, such as neophobia, e.g., responses to novelty, and innovation e.g., problem-solving, particularly in today’s changing world. Bali myna are a critically endangered endemic species, which are a focus of active conservation efforts, including reintroductions. Therefore, gathering cognitive data can aid in improving and developing conservation strategies, like pre-release training and individual selection for release. In 22 captive Bali myna, we tested neophobia (novel object, novel food, control conditions), innovation (bark, cup, lid conditions) and individual repeatability. We found effects of condition and social environment, including longer latencies to touch familiar food in presence than absence of novel items, and between problem-solving tasks, as well as in the presence of conspecifics, compared with being alone, or with conspecifics and competing heterospecifics. Individuals were repeatable in latency responses: 1) temporally in both experiments; 2) contextually in innovation experiment and between both experiments (and approach order), suggesting a stable behaviour trait. These findings are an important starting point for improving conservation strategies in Bali myna and other similarly threatened species.

## Background

Cognitive abilities such as perception, learning, decision making and memory [1], have been long accepted as playing a vital role in animal behaviour in the wild. Crucially, they determine in part an animal’s ability to adapt (i.e., respond flexibly) to variation in ecology and the social environment [2, 3], and individual differences have important fitness and survival implications [4]. Indeed, cognition correlates with fitness measures [5]. As human actions increasingly impact on the environment, animals are often required to adapt quickly to evolutionary novel cues, with new selection pressures placed on cognitive adaptations [6].

Despite this, cognitive research has been largely under-utilised in conservation strategies [3, 6–8]. Understanding how animals perceive and learn about danger and reward can aid conservation, particularly those conservation strategies that rely on manipulating behavioural responses [3, 6]. For example, the way that birds perceive human-made structures, like turbines, has influenced the development of wind farm deterrents [6]. Further, it is not yet known how individual cognitive phenotypes impact survival post-release and establishing this can allow more effective selection of suitable individuals for release.

Reintroduction programmes are a vital conservation tool, but they often fail as released animals cannot forage effectively, recognise predators and/or breed successfully. In animal reintroductions or translocations, the risk of mortality to predation in previously captive animals with little predator experience is high, therefore antipredator training is beginning to be included in pre-release preparations [9]. Antipredator training may be more successful in species that have effective responses to similar predators. For example, New Zealand robins exposed to unfamiliar predators (stoats) have learned to recognise them, impacting on their survival [10]. Recently, there has been a call for closer integration of cognition and conservation research [3, 6, 8]. This type of research is timely and critical given the rapidly declining animal populations worldwide.

Examples of conservation-relevant cognitive abilities include innovation and neophobia. Innovation can be defined as solving a novel problem or finding a different solution to a familiar problem, which influences how animals adjust to new or changing environments [11, 12] (behaviour resulting from a combination of cognitive and non-cognitive processes). For example, innovative common myna (*Acridotheres tristis*) are slower than non-innovative myna to change their behaviour in response to a changing environment [13]. Neophobia – e.g., aversion to novelty - is linked with life-history variation and has fitness implications [4]. Neophobia can aid in avoidance of unfamiliar dangers, though can also prevent adaptation to new environments or foods [14]. How an animal responds to novelty can predict post-release outcomes [15].

An understanding of how species and individuals respond to novelty and approach new problems is vital both for cognitive research and applied conservation, particularly as the world is increasingly urbanised, and many species need to adapt to human-generated environmental changes and the inevitable associated novelty [6]. Individuals or species with higher innovation and lower neophobia may be more adaptable in regard to coping with changing habitat/degradation, though these traits may increase chances of being trapped by humans. Differentiating between responses to these two threats is important as populations/areas face different levels of risk. For example, individual common myna that inhabit urban environments show lower neophobia and utilise novel food resources more quickly compared with those living in rural areas [16].

Furthermore, individuals may show behaviours that are temporally and contextually repeatable, or alternatively, show inconsistency in their responses [17]. This may be influenced by various factors, such as species, task or measures tested, seasonality as well as developmental and social influences [6, 18]. Individual performance may also correlate across tasks. For instance, in feral pigeons (*Columba livia*) and zenaida doves (*Zenaida aurita*), latency to learn a foraging task covaried with individual neophobia level [19].

Bali myna are a critically endangered species that are endemic to Bali. We selected this species because: (1) they are highly threatened (<50 adults in the wild; Birdlife.org); (2) face threats like illegal trapping/ poaching for the pet trade and habitat degradation [20] that could be mitigated through behavioural research and training which must be informed by cognitive research; (3) there is active conservation action with varying success across different sites (need to continually release birds to try to boost small populations and open questions regarding ways to boost survival, such as predator/trapping avoidance and use of novel habitats and safe, novel foods), including reintroduction, which enables pre- and post-release research; (4) there is currently minimal published cognitive/behavioural data on Bali myna though there is a reasonably sized zoo population (~950 individuals across ~170 institutions worldwide, with ~90 individuals in UK zoos; ZIMS, 2021 - zims.species360.org, accessed September 2021).

We aimed to quantify individual and species-level performance in innovation and neophobia tasks in captive Bali myna, using comparable paradigms as in some previous species [13, 14, 21]. We tested (1) innovation through 3 simple problem-solving tasks (flip bark, flip cup and lift lid to obtain preferred insect; 3x 20-minute trials per task) and (2) neophobia through presentation of 3 types of novel objects and novel foods (jelly) placed beside familiar food, compared with familiar food alone (i.e., control; run 3x 20-minute trials per condition for individual repeatability) [17]. Further, we tested whether individual performance correlated across the two experiments, i.e., whether less neophobic individuals were also quicker to approach the problem-solving task(s). We tested individuals within three UK zoos, primarily in social settings as it was not possible to separate individuals for testing. We expected that, similar to other species (e.g., ravens (*Corvus corax*) [22]), social context would influence neophobia and innovation in Bali myna. As the birds were particularly important for the captive breeding programme, and it was necessary for testing to overlap with breeding season due to funding availability, these experimental designs were specifically selected to minimise researcher presence or disturbance in aviaries (e.g., no training requirements).

As there has been little previous published cognitive research with Bali myna, we were reliant primarily on anecdotal/filmed reports before testing. For example, filmed reports of wild-bred juveniles learning to flip cow dung for insects, although their great, great grandparents did not do this in the aviary before release, yet they have worked it out (Donato, 2020, personal communication). These reports indicate innovation may be present in this species, though this has not previously, to our knowledge, been tested. Anecdotal zoo-keeper reports (e.g. Miller, 2021, personal communication) suggested that these birds are relatively neophobic, certainly compared to other similar myna species, like common myna – which are a successful invasive species in Australia and other countries [13]. We expected neophobia to vary across individuals and between conditions, e.g., novel food versus novel object compared with control, as indicated in other species [21, 23]. Finally, we expected that individual performance would correlate across tasks, as in other species (pigeons [24]; corvids [25]; birds and primates [26]). This study provides the first assessment of two key cognitive aspects that influence adaptability in captive Bali myna as a necessary starting point for testing cognition in order to implement findings in active conservation in Bali.

## Methods

We pre-registered this study prior to data collection at OSF: shorturl.at/ftMV1.

### Subjects

Subjects were 22 captive Bali myna (10 males; 10 females; 2 unknown sex) housed across 10 aviaries within three UK zoological collections (Table 1). They were identifiable using coloured or metal leg rings. Subjects were primarily adults (>1 year old, D.O.B. range: 2011-2020), other than two juveniles that were successfully reared by one pair in July 2021. Each zoo housed their birds according to their standard ethical and housing conditions, with a range of aviary sizes, though all (except 1 temporary inside aviary) being primarily outside, with a wide array of perching, planting and substrates available. As it was not possible to individually separate birds at any zoo due to ethical and housing constraints, as well as time restrictions, we tested the birds according to their current housing situation, which included the presence of conspecifics (19/22 subjects) and other bird species (13/22 subjects; Table 1). The other bird species were divided into non-competitors and competitors, based on whether or not they routinely visited the test sites, ate Bali myna food and/or interacted with experimental stimuli/apparatuses (Table 1). Participating in testing was voluntary for the birds – all available birds were present in every trial, other than the two juveniles who were only present for round 2 and 3 (as hatched after round 1 complete). Data collection took place from May-July 2021, which includes the breeding season for this species (timing selected due to funding availability for this limited period). Nest boxes were present in the aviaries that housed male/female pairs for periods of testing, and one pair did successfully reproduce two chicks.

**Table 1.** Subject information.

| UK Zoo | Aviary | Sex<br>(male,<br>female,<br>unsexed) | Age<br>(adult,<br>juvenile) | Mixed exhibit<br>species inc.<br>whether<br>“competitor” or<br>“non-competitor” | Testing<br>site<br>within<br>aviary | Notes |
| --- | --- | --- | --- | --- | --- | --- |
| Birdworld,<br>Farnham | Group<br>(n=7) | 3.4 | Adult | Competitor: 1<br>Lilac-breasted<br>roller, *Coracias*<br>*caudatus*, 3 wonga<br>pigeon,<br>*Leucosarcia*<br>*melanoleuca*, 2 | Main<br>aviary | 1 hatched<br>2018;<br>others<br>2020 |
|  |  |  |  | white-browed robin-chat, *Cossypha heuglini* |  |  |
| Birdworld, Farnham | Pair 1 | 1.1 | Adult | Non-competitor: 2 Edward pheasant, *Lophura edwardsi* | Inside area | Reared 2 chicks in July 2021 – present in aviary during round 2 of testing |
| Birdworld, Farnham | Pair 2 | 1.1 | Adult | None | Covered area of main aviary |  |
| Birdworld, Farnham | Juveniles (n=2) | 0.0.2 | Juvenile | None | Main aviary | Tested with parents for round 2, then alone for round 3 |
| Cotswolds Wildlife Park & Gardens | Pair 1 | 1.1 | Adult | Competitor: 2 white-spotted laughing thrush, *Lanthocincla bieti*, 6 azure-winged magpie, *Cyanopica cyanus*, 2 pink pigeon, *Nesoenas mayeri*, 2 Madagascar partridge, | Main aviary |  |
|  |  |  |  | *Margaroperdix madagarensis* |  |  |
| Cotswolds Wildlife Park & Gardens | Pair 2 | 2.0 | Adult | None in first aviary (round 1 & 2); non-competitor: 1 pink pigeon and two Palawan peacock pheasant, *Polyplectron napoleonis*, in second aviary (round 3) | Main aviary | Moved enclosure July 2021 |
| Waddesdon Manor | Pair 1 | 1.1 | Adult | Non-competitor: 1 Rothchild's peacock pheasant, *Polyplectron inopinatum* | Main aviary |  |
| Waddesdon Manor | Single 1 | 1.0 | Adult | None | Main aviary | Temporary single housing (new arrival) |
| Waddesdon Manor | Single 2 | 0.1 | Adult | None | Main aviary | Temporary single housing (awaiting pairing with new arrival) |
| Waddesdon Manor | Single 3 | 0.1 | Adult | None | Inside house | Temporary single housing (awaiting pairing or relocation) |

### Pilot

Prior to testing, we visited each zoo at least twice to set up test sites (primarily situated where the birds were usually fed), positions for video cameras (min 1 metre, preferably further where possible, from test sites in case birds responded to the camera presence) and recorded latencies to approach familiar food (i.e., regular diet) when fed in the morning (no experimental manipulation).

### Neophobia experiment

#### Apparatus/materials

We included three conditions: control (regular diet); novel food (3cm^3^ blocks of coloured jelly - orange, purple and green); and novel object (human-made items; Figure 1) – each novel item was presented alongside the regular diet (i.e., familiar food). The regular diet was presented in the same food bowl that it would usually be served in at each aviary. Insects (mealworms, waxworms or morio worms) were added to the food bowl. The novel item was typically presented in a familiar food bowl (new bowl present in aviary for several weeks prior to testing). The novel items were selected as such to be comparable with research in corvids [21, 23] so the data may be useful for comparative research [27].

**Figure 1.**
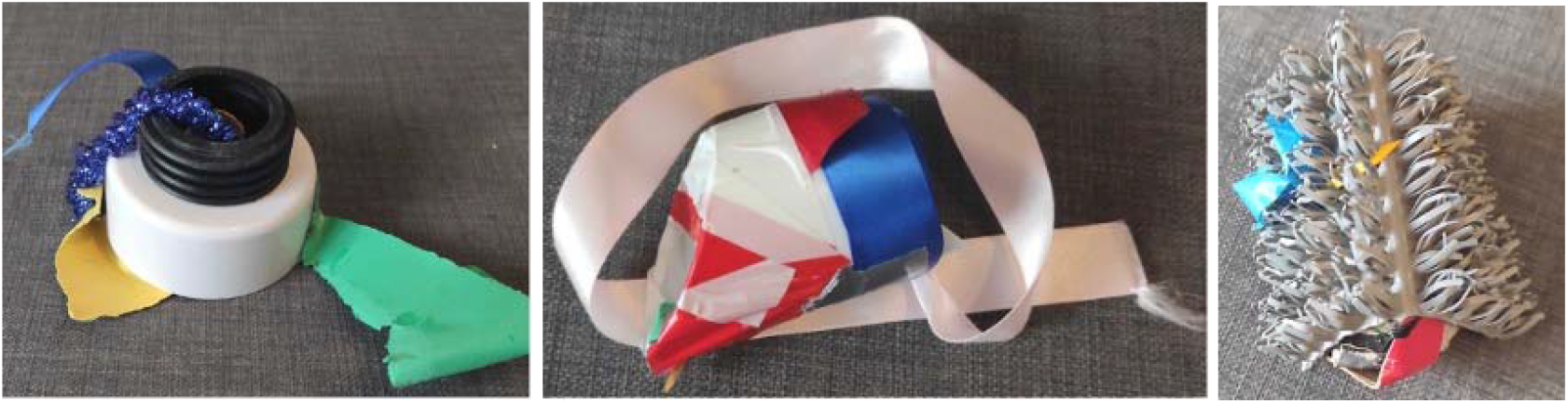
Novel objects.

#### Procedure

We measured behavioural responses to novel items presented alongside familiar food compared with familiar food alone. On novel item condition trials, the novel item was placed ~20 cm from the familiar food bowl, in the same location for each trial so consistent within individual and aviary. For video coding, the trial commenced once the experimenter had left the immediate testing area (i.e., out of camera shot). Each trial lasted 20 minutes in total – determined during piloting to be sufficient time for the majority of individuals to reliably approach the familiar food. Where there was more than one Bali myna subject in an aviary, we established more than one test site using feed sites that already existed or else following at least 2 weeks habituation (i.e., pair-housed aviaries received one or two test sites depending on availability for test sites, the group-housed aviary received three test sites). The experimenter was not present in the aviary during testing.

We ran three tests ‘rounds’ in total – within each round were three trials, one per condition (9 trials total), over 3 days, with approx. 2 weeks between rounds, therefore lasting approx. 6 weeks per zoo. Testing occurred in the morning alongside the daily presentation of their regular diet – therefore the birds were not fed prior to testing, though were not deprived and had access to any leftover food from the previous day as well as any natural foraging opportunities available like wild insects (as all included outside aviary spaces). The control trial (familiar food only) was run on day 2, with the novel food or novel object counterbalanced between day 1 or 3 across aviaries and rounds, so that the control took place within 24 hours of each test condition. The main variable of interest was latency to touch familiar food, indicating the time taken for an individual to touch a familiar food when a novel item was present, with avoidance being interpreted as neophobia (as per [14, 21, 23]).

### Innovation Experiment

#### Apparatus/materials

We included three problem-solving tasks (Figure 2), with a preferred insect as a reward, primarily waxworms or morio worms. Insects were humanely killed by removing their head before testing to prevent them moving away.

**Figure 2.**
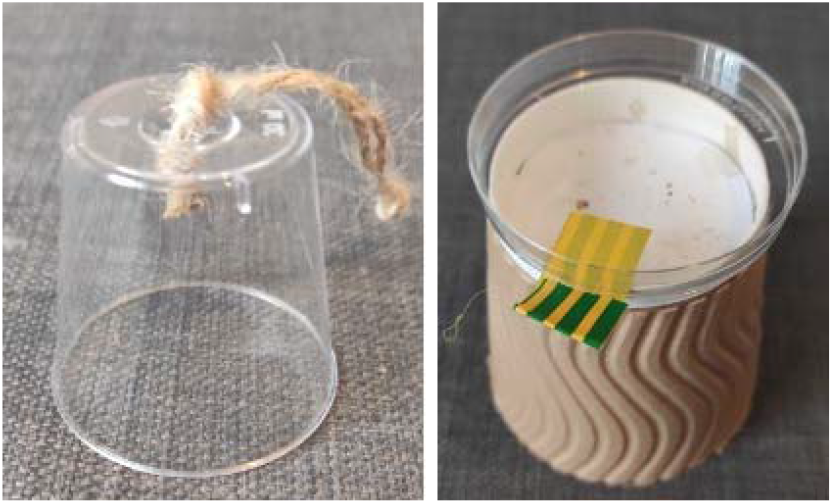
Problem-solving tasks 1 and 2. 1) cup can be lifted to access worm e.g., by pulling string or pushing cup over; 2) lid can be removed e.g., by pushing lid or lifting tab. Problem-solving task 3 was a piece of wood bark that could be pushed or lifted to access worm.

#### Procedure

Each problem-solving task was baited with a preferred reward and required the subject to move an object (lid, cup, bark) to access the reward. In task 1 and 2 (lid and cup), the reward was visible, whilst in task 3 (bark), it was only partially visible (worm placed under bark so the tip of the body was still visible). We selected these tasks as they were relatively simple given that all subjects were unhabituated and unfamiliar with cognitive testing participation, had more than 1 possible method of ‘solving’ and were comparable to previous research with common myna [13]. Each task was presented 3x over 3 days, for 20-minute trials per aviary, over the course of a 6 week period (testing every 2 weeks). Testing occurred in the morning after the neophobia testing was complete. We presented one set of each task per subject for the single- and pair-housed subjects, and four sets for the group-housed subjects. As with neophobia, the experimenter was not present in the aviary during testing, and the video was coded from when the experimenter left the test area(s). If the subject solved the task within the first 5 minutes, the experimenter re-baited it with a new reward item. We measured several variables but namely: latency to touch, and frequency of peck and solve, as well as whether the subject solved the task once or twice per trial.

#### Data Analyses

We recorded all trials and coded all videos using Solomon Coder [28] – the primary coder (E.D.) was unfamiliar with the species and hypotheses prior to coding. We second coded 12% of videos and inter-rater reliability was high: Cohen’s Kappa = 0.8.

For the neophobia experiment, we were interested in two main questions: 1. Testing effects of condition, round and social environment 2. Individual repeatability over round and condition, with the main dependent variable being latency to touch familiar food (0-1200 seconds). Analysis was run using R (version 4.1.0). For Q1, we conducted a Generalised Linear Mixed Model (GLMM) to test whether the main effects of condition (control, novel food, novel object), round (1-3) and social environment (1. alone, 2. conspecific present and/or non-competing heterospecifics i.e., that do not touch Bali myna food, 3. conspecific present and competing heterospecific i.e., that do touch their food) influenced latency to touch familiar food, with individual as a random effect, using likelihood ratio tests (drop1() function) and Tukey comparisons for post-hoc comparisons (package multcomp, function glht()). We checked whether nesting individual within aviary as a random effect, and separately, whether including sex as a main effect improved the model, but did, so we proceeded with individual as a random effect and without sex as a main effect for a more parsimonious model. For Q2, we tested individual repeatability over time (i.e., across rounds) and over condition using intraclass correlation coefficients (ICCs) (per [23]) in SPSS (version 27).

For the innovation experiment, we checked whether frequency to peck correlated with frequency to solve using two-tailed Spearman’s correlations on trials without zeros (73/198 trials) in SPSS. Although 77% of subjects interacted with the tasks at least once, the data were heavily skewed towards zero. We therefore found that GLMM’s were not the most suitable approach, so used non-parametric statistics in SPSS for this analysis – namely, Wilcoxon signed ranks tests and Mann-Whitney U-tests, with Bonferroni corrections applied for multiple comparisons. We compared condition (bark, cup, lid), trial (1-3) and social environment on two variables of interest: 1) latency to approach task, and 2) frequency of peck (which correlated with frequency to solve). We selected these two variables as they showed the highest variability across subjects.

Finally, we tested whether individual performance correlated across the two experiments using intra-class correlation coefficient (ICCs). As subjects were temporally repeatable in both experiments, we created mean scores across round (neophobia) and trial (innovation). We then correlated individual latency to touch familiar food in the object condition of the neophobia experiment with latency to approach in the problem-solving tasks using these mean scores. Furthermore, we used the mean scores to check whether order of approach correlated across the problem-solving tasks and the neophobia tasks within each aviary using ICCs. Example video trials can be found at: https://youtu.be/EngP6mThj4M

## Results

### Neophobia Experiment: Testing Effects of Condition, Round and Social Environment

Latency to touch familiar food differed between conditions (GLMM: *X*^2^ = 62.389, df = 2, p < 0.001) and social environment (*X*^2^ = 8.786, df = 2, p = 0.012), but not between test rounds (*X*^2^ = 3.436, df = 2, p = 0.179). The birds waited longer with a novel object or novel food present compared to the control condition (Tukey contrasts: novel object – control, z = 6.902, p < 0.001; novel food – control, z = 3.665, p < 0.001), and they waited longer when a novel object was present than when a novel food was present (z = 5.015, p < 0.001) (Figure 3A). They waited longer when conspecifics and non-competing heterospecifics present (2) compared with when conspecifics and competing heterospecifics present (3; Tukey contrasts: z = −2.961, p = 0.008). There was no difference in latencies when alone (1) compared to conspecific and/or non-competing heterospecifics (2; z = 1.106, p = 0.505), or alone (1) compared to conspecific and competing heterospecifics present (3; z = −0.952, p = 0.603; Figure 3B). Subjects touched the novel food in 3 of 62 trials (4.8%) and novel object in 0 trials, therefore latency to touch the novel items was not a useful measure for testing.

**Figure 3.**
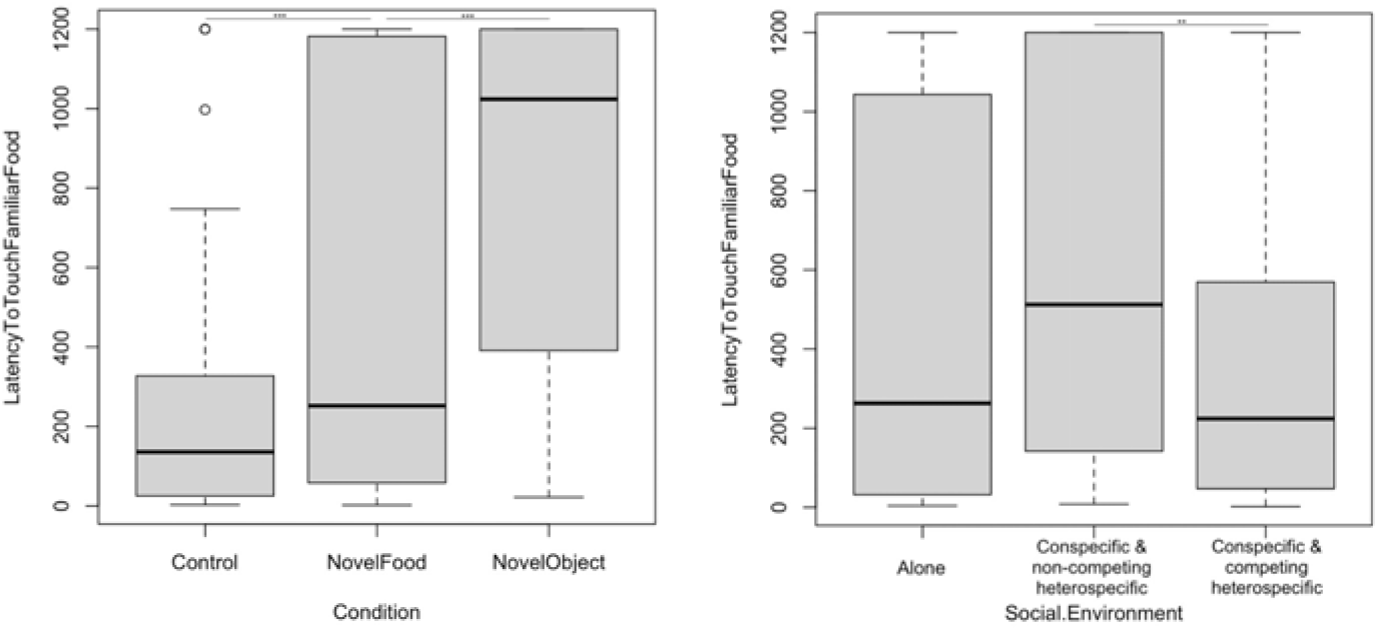
Latency to touch familiar food (seconds) differed by A) condition and B) social environment. Raw data; lines represent median. *** p < 0.001; ** p < 0.01

### Neophobia Experiment: Individual Temporal and Contextual Repeatability

In the neophobia experiment, we found that individuals were temporally repeatable across 3 test rounds (intra-class correlation coefficient: N = 22, ICC = 0.632, p < 0.001, CI = 0.435-0.768). Individuals were not contextually repeatable across novel item conditions (novel object, novel food) in their responses to novelty (ICC: N = 22, ICC = 0.278, p = 0.103, CI = - 0.199-0.565). Within condition, they were temporally repeatable within the control condition, but not within the two novel item conditions (control: N = 22, ICC = 0.0.543, p < 0.02, CI = 0.038-0.805; novel object: N = 22, ICC = 0.287, p = 0.182, CI = −0.501-0.696; novel food: N = 22, ICC = 0.278, p = 0.183, CI = −0.521-0.692).

### Innovation Experiment: Testing Effects of Condition and Social Environment

17 of 22 (77%) subjects approached and solved at least one trial/task. Frequency to peck correlated with frequency to solve (Spearman’s correlation: trials with zeros removed: r(20) = 0.302, p = 0.01). Latency to approach and frequency of pecking problem-solving tasks differed across conditions, as subjects waited longer to approach and pecked less frequently in the bark than cup condition (Wilcoxon signed ranks test: latency to approach - Z = 0.475, p = 0.028; frequency of peck – Z = −0.458, p = 0.036), with no difference between cup and lid (latency - Z = −0.5, p > 0.999; frequency – Z = 0.142, p > 0.999), or bark and lid tasks (latency - Z = 0.425 p=0.06; frequency – Z = −0.317, p = 0.249).

Latency to approach and frequency of pecking also differed across social environments. Specifically, subjects waited longer to approach when with conspecifics and non-competing heterospecifics present (2) compared with when alone (1) or when conspecifics and competing heterospecifics were present (3) (Mann-Whitney U test: 1 vs 2 – U = −33.414, p = 0.011; 2 vs 3 U = 30.315, p = 0.001; 1 vs 3 U = −3.099, p > 0.999; Range = 0-1200 seconds; Mean = 718.4; Figure 4A). Subjects also pecked less when conspecifics and non-competing heterospecifics present (2) compared with conspecifics and competing heterospecifics present (3), with no difference compared to being alone (1) (Mann-Whitney U test: 1 vs 2 – U = 20.833, p =0.147; 2 vs 3 U = −20.357, p = 0.019; 1 vs 3 U = 0.475, p > 0.999; Range 0-21 pecks; Mean = 1.4; Figure 4B).

**Figure 4.**
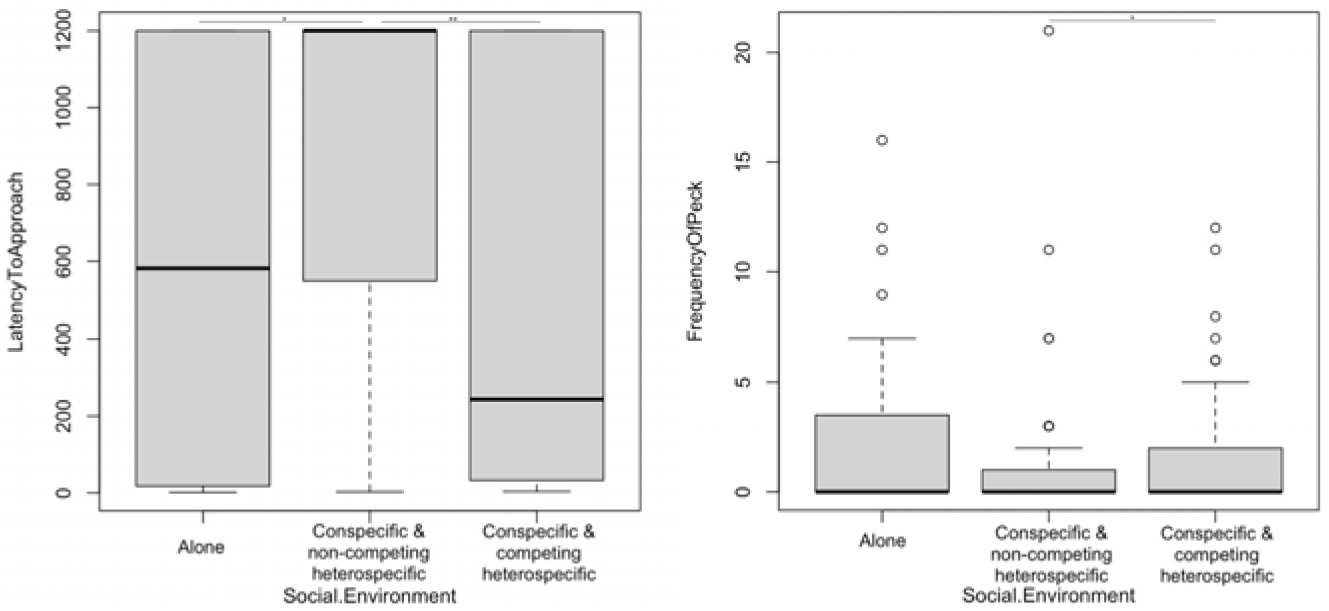
Social environment effect on A) latency to approach (seconds) and B) frequency of peck on problem-solving tasks. Raw data; lines represent median. *** p < 0.001; ** p < 0.01; * p < 0.05

### Innovation Experiment: Individual Temporal and Contextual Repeatability

Individuals were temporally repeatable (across 1-3 trials: ICC = 0.547, p < 0.001, CI = 0.313-0.710) and contextually repeatable in latency to approach the problem-solving tasks (across bark, cup, lid conditions: ICC = 0.317, p = 0.040, CI = −0.048-0.570).

### Individual-level Performance across Both Experiments

Using a mean score across round/trial, individual latency to approach three problem-solving tasks correlated with latency to touch familiar food in presence of novel object (n=20, ICC = 0.763, p < 0.001, CI = 0.533-0.896). Using the mean score, the order of approach within aviary correlated across the three problem solving tasks and the object neophobia condition (n=17, ICC = 0.915, p < 0.001, CI = 0.823-0.966; note. 3 subjects tested alone and 2 subjects – the 2 juveniles - not tested in the bark condition so excluded from analysis).

## Discussion

We tested neophobia (latency to touch familiar food in presence of novel object or novel food) and innovation (latency to approach and frequency of pecking three simple problem-solving tasks) in captive Bali myna. We found effects of condition (neophobia: control, novel object, novel food; innovation: bark, lid, cup) and social environment (alone, presence of conspecifics and/or heterospecifics that do or do not touch food, i.e., competitors or non-competitors) on both neophobia and innovation. Individuals were temporally repeatable, though not contextually repeatable in their responses to novelty, while being temporally and contextually repeatable in responses to the problem-solving tasks. Individuals also showed repeatability in their latency responses and order of approaches across both experiments. These findings indicate that, for example, an individual that is quick to touch familiar food beside a novel object is also quick to approach a problem-solving task, and subjects within each aviary are likely to approach the task in a similar order across trials. This study is one of the first to test cognition in Bali myna and provides support for the feasibility of this study species for future research.

Our findings indicating individual repeatability suggest that behavioural responses to novel objects and foods, as well as simple problem-solving foraging-based tasks, may reflect stable traits in this species. Social context has been shown with other species to either facilitate or inhibit behaviours, including neophobia and exploration [18, 29, 30]. For instance, observing group-members eating familiar food facilitates acceptance of novel foods in capuchin monkeys [31]. Bali myna behaviour was influenced by presence of others in both experiments. It appears that the specific identities and/or behaviour of others present played a role, given that conspecific (typically – though not always - a partner) and/or non-competing heterospecifics tended to inhibit Bali myna behaviour, whereas presence of conspecifics (such as the group) and competing heterospecifics (routinely interacted/ate at Bali myna food sites/stimuli) tended to facilitate Bali myna responses. This finding may reflect a ‘socially-induced’ neophobia, where individuals wait for others to take the risk of approaching first, or alternatively related to rank, where they have to wait for access (Mainwaring et al., 2011). The latter finding may be likely with the Bali myna, given they show a consistent order of approach within aviaries which may relate to rank. The importance of the relationship and/or identity of conspecifics has also been shown to influence exploration and/or neophobia in corvids, wolves and dogs [18, 22, 30]. The tested group of young Bali myna in particular presents a rare opportunity (given that this species is most often held in pairs) for future social-based experiments, such as facilitation and tolerance around food sources [32].

The problem-solving tasks selected were similar to one another and simple – lifting or pecking at an object to obtain a visible reward underneath. Despite this, we found differences in responses across conditions. The longer latencies for the bark over the lid and cup conditions is likely due to this being the first task that was tested (i.e., test round 1).

Alternatively, it may be related to the reward (worm) being less visible under the bark than inside the transparent cup or lid. Future work may explore understanding of object permanence to test whether reward visibility influences behavioural responses in problem-solving tasks. We selected the frequency of peck, rather than rate of solving, measure as pecking and solving correlated with one another, and pecking showed higher variability.

The main study limitations were uncontrollable aspects of the testing environments – namely variable presence of conspecifics and heterospecifics, which we included as a factor in the analysis. Some heterospecifics had little recordable impact on Bali myna interactions with food or experimental stimuli (e.g., ground-dwelling species like pheasants) thus were referred to as “non-competitors”, while others (e.g., spotted laughing-thrush) routinely interacted with these items thus were “competitors”. Interestingly, despite appearing to be quite neophobic (i.e., stronger reaction to novel items than control, particularly to novel objects), the Bali myna anecdotally frequently appeared to be one of the more dominant species in mixed-species aviaries as they displaced others (e.g., azure-winged magpies) from test/food sites. We were restricted in timing of data collection due to funding availability therefore testing overlapped with breeding season, which may impact on cognitive performance, motivation and participation. Indeed, one pair did successfully reproduce during testing, which provided a unique opportunity to test two Bali myna juveniles shortly after fledging in the presence of the parents, as well as while alone.

These were captive zoo-housed individuals limiting generalisation across the species. Future work should aim to include A) larger captive sample size generally and B) wild/ reintroduced birds. Neophobia and innovation could be tested further using different tasks, such as novel predators, variety of novel foods, and more complex problems-solving tasks. Similarly, as neophobia has been found to be context-specific in other species (e.g., corvids [33, 34]), it would be useful to explore the flexibility and manipulations of this behavioural response to novelty. For instance, increasing (e.g., via pairing with aversive stimuli) neophobic reactions to dangerous items, like traps, or decreasing (e.g., via habituation) neophobic responses to novel safe foods prior to release. Other cognitive aspects that are relevant to adaptability, such as social learning i.e., learning from others, would also be useful to test for applying to conservation actions. For example, social facilitation during foraging (capuchin monkeys [35]; carrion crows [32]; bats [36]) and exploring the link between different cognitive abilities, like innovation and social learning [24, 25]. Our present finding that Bali myna are influenced by social context indicates that this would be a useful avenue for future work.

## Conclusion

We tested two conservation-relevant cognitive abilities in a little-studied, critically endangered bird species, which could be further implemented across other species, for instance through the ManyBirds framework [27] and utilised in applied sciences. Additionally, behavioural research contributes to conservation by encouraging positive public perception [6], which is particularly important for preventing poaching for the pet trade - a major threat to Bali myna and other species. These findings are promising for the potential of future research with Bali myna and similarly threatened species, particularly those that may be available for both captive and fieldwork, with active conservation programmes including reintroductions.

## Acknowledgements

Thank you very much to our participating UK zoological collections: Waddesdon Manor (National Trust/ Rothschild Foundation), Cotswolds Wildlife Park and Gardens, and Birdworld. Special thanks to: Ian Edmans, Gavin Harrison, Llyr Davies (Waddesdon Manor), to Helen Hitchman, Chris Green, Richard Wardle, Natalie Horner (Cotswolds Wildlife Park and Gardens), and to Duncan Bolton, Kat Nicola, Polly Bramham, Natalie Marshall, Rebecca Ive and Ellie Wiczling (Birdworld) for facilitating this research. Thank you to Megan Lambert for helpful feedback on a manuscript draft.

## Ethics Statement

For animal research, all applicable international, national and/or institutional guidelines for the care and use of animals were followed. This non-invasive behavioural study with birds was conducted adhering to UK laws and regulations and was covered under a non-regulated procedure through University of Cambridge, approved by the Home Office appointed Named Animal Care and Welfare Officer, Named Veterinary Surgeon and Chairperson for the Psychology and Zoology Department Animal User’s Management Committee.

## Data Accessibility

The full dataset and R script are available at Figshare: DOI 10.6084/m9.figshare.16974298.

## Author Contributions

R.M. conceived the study idea, research design, project managed the study, analysed the data, produced the figures, and was awarded funding to support the study. E.G.P. contributed to the research design. R.M. and E.G.P. collected the data. E.G.P. and E.D. coded the videos. R.M. wrote the manuscript, with E.G.P. and E.D. providing feedback on the manuscript.

